# Single unit electrophysiology recordings and computational modeling can predict octopus arm movement

**DOI:** 10.1101/2024.09.13.612676

**Authors:** Nitish Satya Sai Gedela, Sachin Salim, Ryan D. Radawiec, Julianna Richie, Cynthia Chestek, Anne Draelos, Galit Pelled

## Abstract

The octopus simplified nervous system holds the potential to reveal principles of motor circuits and improve brain-machine interface devices through computational modeling with machine learning and statistical analysis. Here, an array of carbon electrodes providing single-unit electrophysiology recordings were implanted into the octopus anterior nerve cord. The number of spikes and arm movements in response to stimulation at different locations along the arm were recorded. We observed that the number of spikes occurring within the first 100ms after stimulation were predictive of the resultant movement response. Computational models showed that temporal electrophysiological features could be used to predict whether an arm movement occurred with 88.64% confidence, and if it was a lateral arm movement or a grasping motion with 75.45% confidence. Both supervised and unsupervised methods were applied to gain streaming measurements of octopus arm movements and how their motor circuitry produces rich movement types in real time. Deep learning models and unsupervised dimension reduction identified a consistent set of features that could be used to distinguish different types of arm movements. These models generated predictions for how to evoke a particular, complex movement in an orchestrated sequence for an individual motor circuit.

## Introduction

The octopus has many features that makes it advantageous for pursuing a holistic understanding of movement, cognition and behavior: It has a highly developed nervous system containing 500 million neurons and a large brain ^1–3^; Each of the eight arms contains an axial nerve cord (ANC) which resembles and acts like the vertebrate’s spinal cord; Hundreds of suckers and sucker ganglia act as the peripheral nervous system and can demonstrate a large repertoire of behaviors ^4,5^; it has distributed control of its nervous system. One of the advantages of using this animal model, is that an electric or tactile stimulation to the octopus’s denervated arm, can still trigger movement that is similar in kinematics to movement of an intact arm as was previously demonstrated ^6–9^. This suggests that the octopus has a simplified neural program embedded within the arm itself and is adaptable to various degrees of input from visual, sensory and motor brain areas ^6,10,11^. The octopus’s simplified nervous system holds the potential to use machine learning and advanced statistical analysis techniques to develop computational models that could predict motor behavior.

Recently, there has been growing interest in recording electrophysiology signals from octopus nervous system: single unit and intracellular recordings from slices obtained from octopus had revealed principles of learning and memory ^12^, and multi-unit and local-filed potentials were recorded from octopus arms ^11^ and the nerve ring responsible for arm coordination ^13^. Recent developments have also shown the capabilities of recording brain signals from awake octopuses ^14^. These studies revealed fundamental concepts regarding the octopus nervous system, motor control and coordinated movement. However, thus far, the electrophysiology recordings have only consisted of continuous local field and electroencephalogram data. For this study, as described below, we instead used very small diameter carbon fiber electrodes for unit recording. Carbon fiber electrodes are strong enough to penetrate tough neural tissue when sharpened ^15,16^.They also do very little damage to the neurons of interest ^17^. They can be sharpened in a way that preserves a small electrode surface area, enabling high amplitude spikes ^18^.

Understanding the trajectory and dynamics of arm movement is crucial to develop neuroprosthetic devices and robotic limbs that will allow reaching and grasping. Current Brain Machine Interface (BMI) systems are based on decoding algorithms that use the neural signals to control the external device ^19,20^. However, these devices do not provide enough independent degrees of freedom of the arm, and usually control even simple motions of the lower limbs. Recent studies using different computational techniques including machine learning (ML) algorithms and Artificial Intelligence (AI) have shown the ability to predict several aspects of arm reaching from electrophysiology data ^21–23^. To improve future BMI devices, it will be crucial to further reveal the neural mechanisms behind how diverse movements are represented in the measured electrophysiological signals and how these representations relate to distinct kinematic features of the behavioral response (position, velocity, muscle activity, direction, and more) [6-10]. Octopus movement research can inspire new development for flexible and independent neuroprosthetic limbs.

While the octopus demonstrates useful complex and flexible movements, these kinematics must first be measured and quantified for correlation with electrophysiological signals. Analyzing movements in an automated manner can quickly provide crucial insights into animal behavior that would otherwise be too time-intensive or costly to manually characterize. Rather than hand-labeling the position of an animal in a tank, for instance, computer vision tools can be used to automatically report its x and y spatial locations in an image, which could subsequently be correlated with neural firing to identify location-sensitive cells. In many situations, however, a pre-defined quantity such as spatial location may not be the best metric for characterizing behavior. More complex and ethologically relevant behaviors, such as exploration, reaching, or grasping, are better defined by their motion with respect to the animal’s body or to the sequence in which they are performed ^24^. Thus, methods are needed for automated position extraction, pose estimation, and behavioral feature identification.

Both supervised and unsupervised machine learning methods can be used across a wide variety of animal models to classify and cluster behavioral features into these more relevant phenotypes ^25–27^. Supervised machine learning effectively quantifies behavioral data, while unsupervised clustering objectively uncovers inherent structures within datasets, aiding in identifying continuous movement spaces or distinct movement type clusters. While characterizing behavior itself is important for understanding animals and their nervous systems, it lacks the ability to test hypothesized brain-behavior links. That analysis requires models that are able to correlate neural activity and behavioral responses to perturbations. By learning behavioral features associated with various experimental paradigms, we could then correlate what aspects of the environment or stimuli are significantly driving these behaviors. Identifying a set of movements and their orchestrated sequences empowers the construction of simplified yet accurate representations for a particular task, shedding light on underlying mechanisms of e.g., the motor circuits involved in reaching. Finally, with the development of tools that allow real-time analysis with minimal latency ^28–30^, we can also consider closed-loop experimental paradigms that adapt stimulation parameters based on instantaneous behavioral responses. With immediate analysis of how different behavioral features vary during neural stimulation, we could construct models that learn how best to evoke a particular, complex movement in an orchestrated sequence for a particular motor circuit.

This type of data-driven approach could unveil individual behavioral motifs, control circuits, and ultimately contribute to advancements in more flexible and adaptable prosthetics; notably in goal-oriented grasping movements for individuals with limb loss or spinal cord injuries. Here, using state-of-the-art carbon fiber arrays that provided single-unit electrophysiology recording capabilities with ANC neurons ^18^, we obtained simultaneous video recordings, neural patterns and arm kinematics. To trigger movement, descending stimulation was delivered directly on the ANC, and ascending tactile stimulation was delivered to the base of the arm, close to the electrodes, and to more distal portion of the arm. Machine learning models were build to then predict resultant octopus arm movement and learn what specific behaviors could be decoded to the original stimulation.

## Methods

### Experiments & data acquisition

All procedures were approved by the Institutional Animal Care and Use Committee at Michigan State University. Adult *Octopus bimaculoides* (n=7) were anesthetized according to protocols published by Butler-Struben et al. ^31^. From each animal we recorded from two arms (L2 and R3), and the recordings of each of the arms was spaced at least three weeks apart. Once the arm was removed, and the proximal end of the arm was restrained in a tray that was continuously perfused with filtered saltwater. The muscles at the base of the arm were dissected, revealing the ANC. High-density carbon fiber of 16 electrodes array was inserted transversely into the exposed ANC (**Figure 1**).

**Figure 1:**
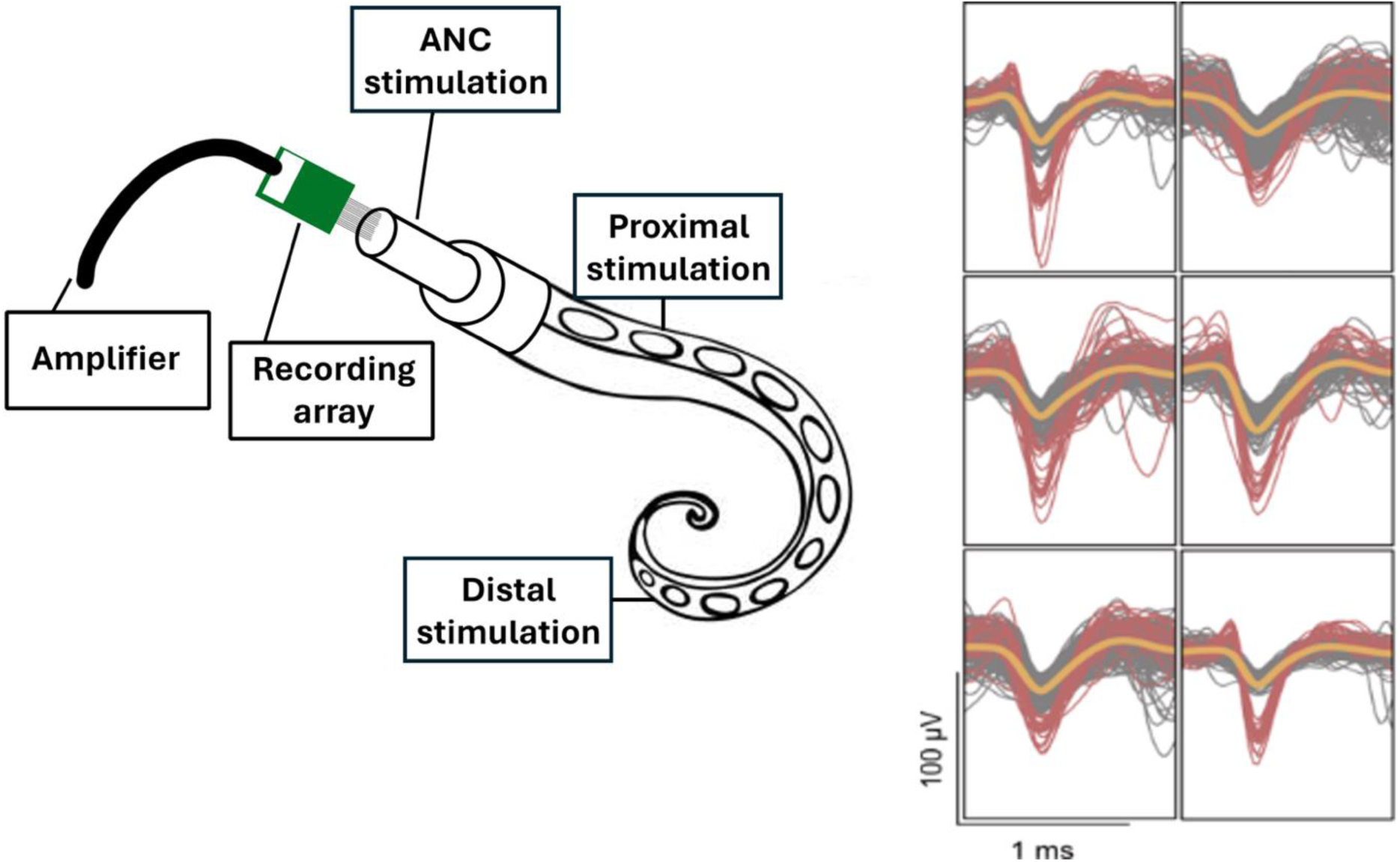
The experiment setup. The 16-electrode array was inserted into the exposed axial nerve cord (ANC). Tactile and electrical stimulation were delivered into three different locations: directly onto the ANC, proximal, and distal to the electrodes’ placements. The carbon electrode array recorded single unit potentials. Representative single units are shown on the right.

To determine if there was a difference in stimulation response, mechanical (tactile) stimulation was performed using plastic forceps and an applied force of 2-3lbf. Electrical stimulation was performed using a single electrode that delivered 5 mA, 100Hz pulses for 50ms. The arm was stimulated in three different locations: directly on the exposed ANC to reflect efferent stimulation, and on regions proximal and distal to the electrode placement, reflecting afferent stimulation. Statistical analysis of the observed distribution of observed movement responses showed that no feature could be used to significantly differentiate between electrical and mechanical stimulation cases across all three stimulus locations. Therefore, all stimulation trials in each of the locations were grouped.

Intan Recording System (Intan Technologies, Los Angeles, CA) and Spike2 (Cambridge Electronic Design, Cambridge, UK) were used to record signals. Spike2 was used for initial signal processing. The signals were filtered with a bandpass second order Butterworth 0.1 to 3kHz and the threshold for action potential (AP) detection was set at 4 standard deviations ^32–34^. APs had an average duration of 1.44ms. A total of 95 experiments resulted in 1520 traces of recordings. For modeling analysis, recordings from three electrodes in the array that showed the highest activity were selected (275 traces).

For movement recording, a webcam was positioned over the recording chamber and videos were recorded simultaneously with electrophysiology recordings via the Spike2 interface. Movement was first manually classified into 3 distinct categories: no movement (NM, “0”), lateral movement (LM, “1”), and a curl (CM, “2”).

Electrophysiology data of the entire 1520-unit recordings traces was processed using Plexon (Plexon Inc, Dalas, TX) offline sorting software with 250 Hz Butterworth filters applied to the raw data. Spikes that had passed the threshold were identified as “units” in the analysis. The processed data was analyzed using Python. The Python script extracted the number of spikes for each 50ms time bin of the 16 channels and summed them. Prism software (Graphpad software Inc, San Diego, CA) was used for statistics. Outlier data was filtered using the 1% ROUT method. ANOVA (Brown-Forsythe) test was used to assess the statistical significance of the relationships between spikes and movement, spikes and stimulation location, and location of stimulation and type of movement. Statistically significant results were considered to be p<0.05.

### Modeling to predict movement from neural activity

We utilized the One Hot Encoding (OHE) technique ^35,36^ to convert the categorical feature into binary features. The OHE technique converts a categorical feature into multiple binary features, where the number of binary features is equivalent to the number of distinct categories in the original categorical feature. This technique assigns a value of 1 to the binary feature corresponding to the specific category for each instance/sample and a value of 0 to all other binary features.

We utilized Cramer’s V to understand the significance of each OHE feature with the categorical target. Cramer’s V is a technique used to measure the degree of association between two categorical features. This technique is based on the chi-square statistic test, and Cramer’s V value ranges from 0 to 1, where 0 indicates no association between the variables, and 1 indicates a perfect association between the variables.

An additional machine learning based method, the Feature importance analysis ^37,38^ can reveal the degree of importance of all features, including categorical and discrete, binned, features to predict the target. The analysis was conducted on both Binary-class (movement/no movement) and Multi-class (no movement/movement/movement with a curl) datasets to identify which features were most influential in predicting the movement outcomes. This analysis was essential to understand the underlying factors contributing to the model’s predictions, optimize the model’s performance, and provide insights into the key drivers of movement patterns.

Overall, an array of 16 different machine learning models was trained on our rich datasets, demonstrating a comprehensive application of diverse machine learning techniques across several categories. Tree-based models like the Decision Tree, ensemble techniques such as robust methods like Random Forest and Extra Trees Classifier, as well as powerful boosting approaches such as Extreme Gradient Boosting (XGBoost), Light Gradient Boosting Machine (LightGBM), Gradient Boosting Classifier, Adaptive Boosting (Ada Boost), and the state-of-the-art CatBoost were employed. Advanced classifiers like the Ridge Classifier, which is based on linear regression techniques but includes regularization, and SVM with a Linear Kernel, which excels in high-dimensional spaces, were also employed. Additionally, simpler yet essential models like the Dummy Classifier were used to establish baseline performances. Our model set further incorporated classical statistical methods, including Logistic Regression and Naive Bayes, alongside discriminant analysis techniques like Linear and Quadratic Discriminant Analysis. The array also included instance-based learning methods like K-Nearest Neighbors (KNN). This varied and methodologically rich collection of models ensured rigorous and nuanced analysis, providing robust and detailed insights.

### Modeling to identify stimulation from resultant behavior

To track the motion of the octopus arm, we first employed DeepLabCut (DLC) for markerless keypoint tracking and pose estimation ^28,30^. This widely-used software package utilizes deep neural networks and transfer learning to achieve accurate 2D and 3D markerless pose estimation for defining and tracking specific points of interest. Out of the total 234 videos, 16 videos with different camera angles, applied stimuli, and observed motion types were selected to train our octopus-specific model. 16 images from each of these videos were then selected by DLC as representative and diverse samples of the octopus arm’s movement, as determined by k-means clustering. Images that were blurred and where the octopus was heavily obscured were then manually dropped from the training set (typically 0-4 images per video). Finally, the images were hand-annotated to label 17 (approximately) equidistant keypoints along the arm using a GUI provided by the DLC package (**Figure 1A**).

Next, we took the ResNet-50 model supplied by DLC, pre-trained on the large and well-established ImageNet dataset, and further trained it using our annotated frames of the octopus arm. This training was done on a lab workstation with a single GPU (NVIDIA GeForce RTX 4070 Ti) and took 3.5 hours to run 150,000 iterations. Once the training was complete, the final model was employed for real-time keypoint tracking of all videos in the dataset. The model output reported x and y location predictions for each of the 17 keypoints in each frame, accompanied by an associated prediction likelihood value.

To comprehensively quantify the entire arm’s motion, a range of significant kinematic features were computed from the x and y predictions. Specifically, we defined θ as the angle formed by the proximal and distal segments (the angle between the stationary base and the tip of the arm), and its instantaneous angular speed as a difference of θ across consecutive frames scaled by frame-per-second to convert to SI units. We also considered the absolute speed of each keypoint v, later focusing on just the distal point as significant. To provide an initial quantification across time, the mean and maximum values of the above features were calculated over three distinct non-overlapping time intervals after stimulation: 0-1 second, 1-2 seconds, and 2 or more seconds (**Figure 2b**). These defined features can be well understood, linked as they are to specific locations along the arm. For example, the distribution of the maximum angular velocity of the arm in the 0-1 second time period post-stimulation is clearly different for the videos hand-labeled as having ‘No movement’, ‘Movement’, and ‘Movement with arm curl’ (**Figure 2c**), as might be intuitively expected.

**Figure 2:**
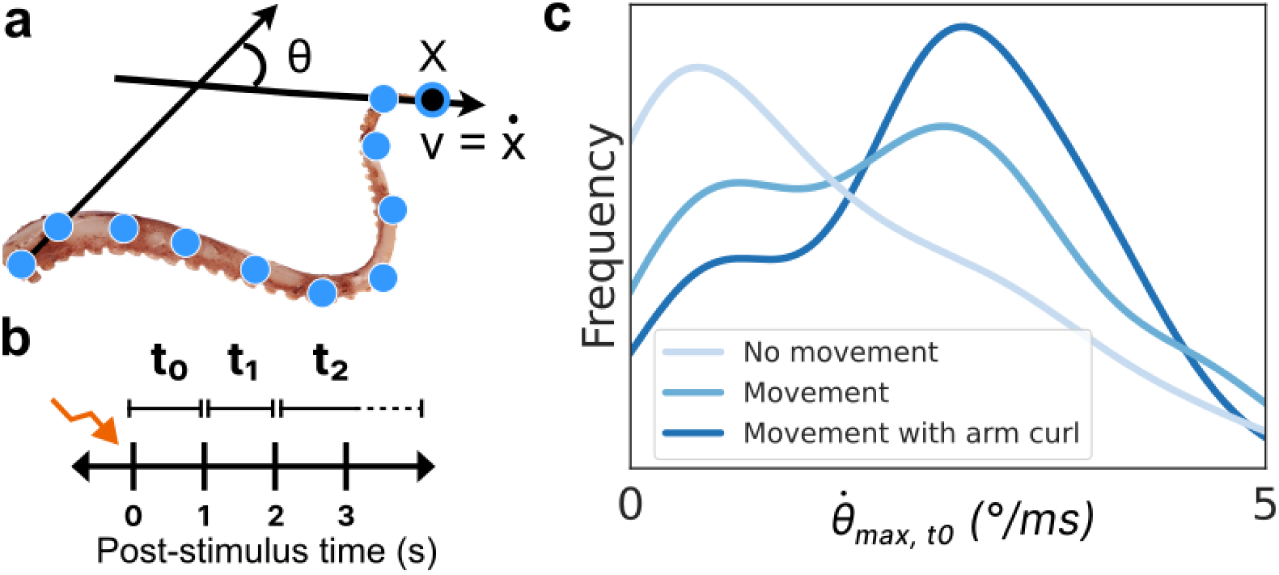
Keypoint tracking and feature extraction. a, 17 roughly equidistant points were hand-labeled along the length of the arm. Using these keypoint positions, various metrics were computed for subsequent analysis including the overall angle made between the (stationary) base and the tip (θ) and the angular and keypoint velocities. The distal-most keypoint (x) and its velocity (v) were found to be significant in distinguishing motion types. b, To quantify motion across time, three intervals post-stimulation were considered: the first (t0) and second (t1) seconds, where most motion occurred, and 2 or more seconds (t2) until any observed motion ceased. c, The histogram of one example metric, the maximum angular velocity in t0, is plotted (using a kernel density estimator) for each of the human-labeled movement categories. As expected, the ‘No movement’ videos have very low or zero angular velocity, whereas ‘Movement with arm curl’ videos tend to have higher maximum angular velocity.

We then employed a second method of analysis that did not rely on keypoints, as typical keypoint tools rely on obvious features such as joints or consistent markings that the octopus arm lacks. We used an unsupervised streaming dimension reduction algorithm known as Procrustean SVD (proSVD) ^29^ to identify features within the videos and how they varied across time, without any pre-training or knowledge of what the videos contained. Unlike conventional SVD methods, proSVD stands out by ensuring the selection of a stable feature set across time, offering dependable results even in the initial phases of data collection. We reduced the videos to 4 bases, or features, and quantified the discovered motion with the L2-norm of each basis vector. Additionally, to optimize processing efficiency, specialized code was developed to crop the videos precisely around the identified DLC keypoints with a 20 pixel margin before handing the cropped videos to proSVD. This tailored step proved instrumental in eliminating superfluous background elements (anything not an octopus arm), which significantly sped up subsequent processing stages.

## Results

### Single and multiunit analysis

Carbon fiber electrodes successfully recorded single and multiunit activity from the ANC, as shown in Figure 1. The total number of units in the first 50ms and 100ms after stimulation were calculated for each of the 16 channels, which resulted in 1520 traces. The movement was based on video analysis and was classified into 3 distinct responses: no movement (“NM”), lateral movement (“LM”), and a movement that consisted an arm curl (“CM”). **Figure 3** shows the number of units in each movement. To test if the number of units occurring immediately after stimulation is different for each movement response, an ANOVA analysis was performed. Results showed that there was a significant difference between the groups means for each movement response type (F(2.00, 42.09)=4.10, p=0.023)). There was also a significant difference between the number of units occurring 50-100ms after stimulation (F(2.00, 46.37)=7.36, p=0.0017). In both time frames, the lateral movement showed the greatest number of units activity. This may suggest that an arm curl is a reflexive response which requires less neural activation.

**Figure 3.**
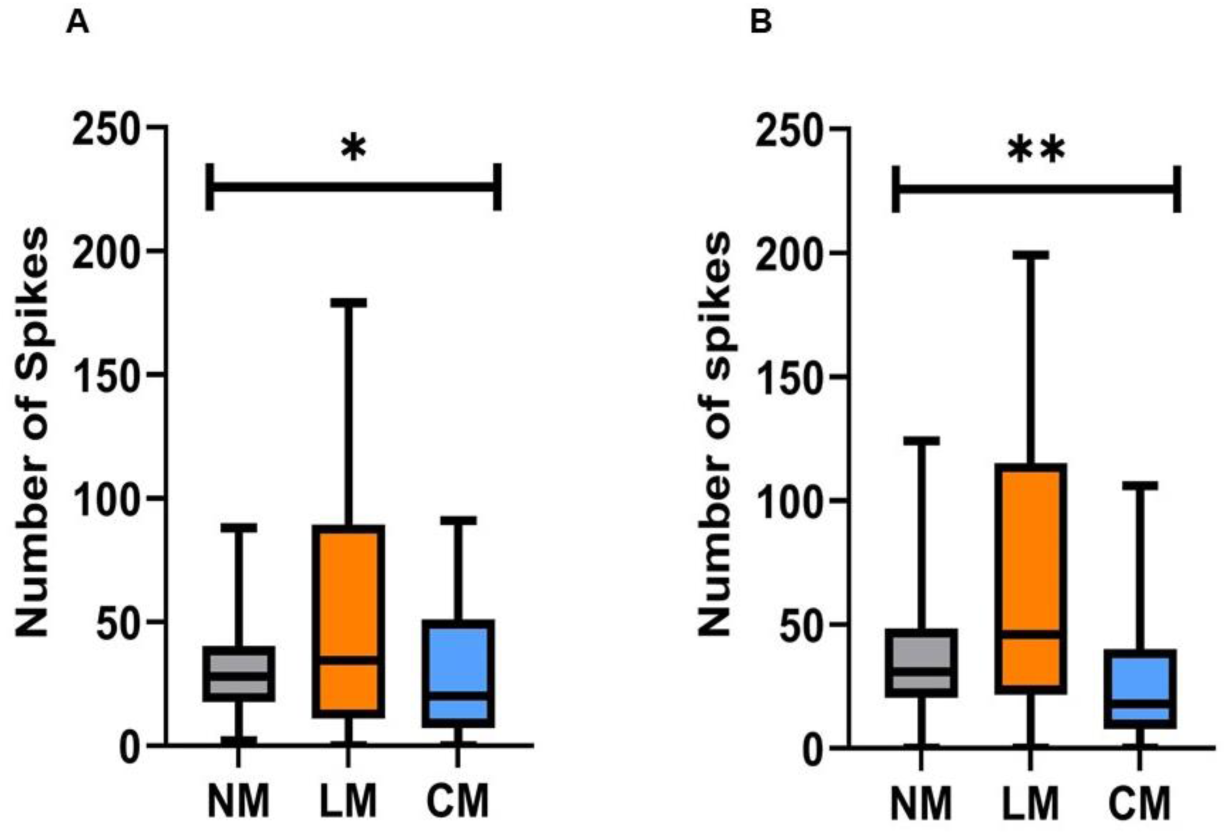
The number of units and movement. (A) The average number of unit responses within 50ms after stimulation evoked different movement response. (B) The average number of unit responses within 50-100ms after stimulation and the evoked movement response. In both time periods, the greatest number of units was found to be associated with lateral movement (LM). (No movement (MN), movement with an arm curl (CM); ANOVA, *<0.05; **<0.005; n=7 octopuses, data obtained form 14 arms).

We then tested if there was a difference between the number of units occurring as a response to the location of the stimulation. The results demonstrate in **Figure 4** show that in the first 50ms and 100 ms after stimulation there is a significant difference between the groups means (F(2.00, 54.23)=6.062, p=0.0042) and (F(2.00, 68.75)=4.72, p=0.012), respectively. In both time frames, the Cord stimulation showed the least variance in the number of units, compared to Distal and Proximal arm stimulation, suggesting that afferent stimulation results in a consistent response. Results also show that the number of units evoked by a Distal stimulation significantly increases over time which may suggest a mechanism to amplify distant signals (paired T-test analysis, p=0.0067). The number of units between the first 50ms and 100ms did not change in response to Proximal or Cord stimulation.

**Figure 4.**
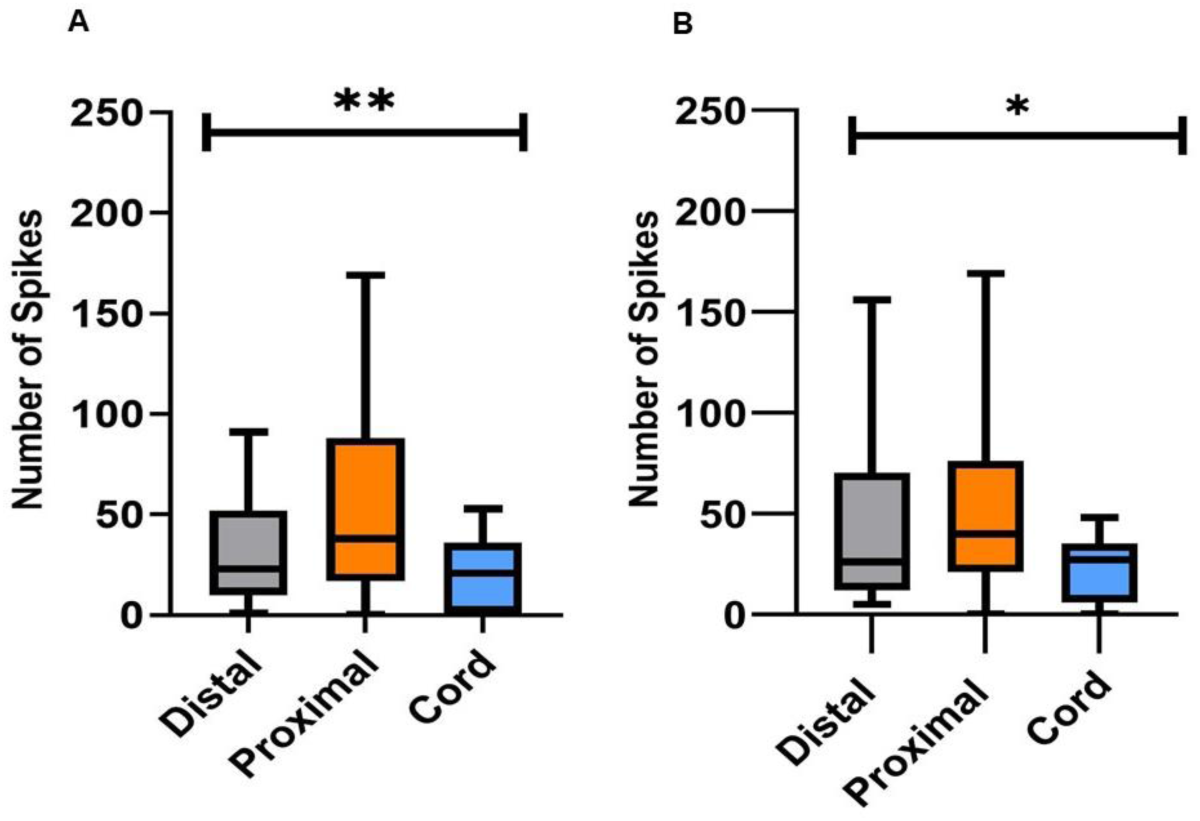
The stimulation type and number of spikes. (A) The average number of unit responses within 50ms after stimulation and the location of the stimulation. (B) The average number of unit responses within 50-100ms after stimulation and the location of the stimulation. Between the two time periods, the greatest increase in units was found to be associated with distal stimulations. (ANOVA, *<0.05; **<0.005; n=7 octopuses, data obtained form 14 arms).

We then sought to determine the probability of the type of stimulation to evoke a specific movement response. **Figure 5** shows the probability of movement response given the type of stimulation. Distal stimulation showed a clear preference to induce movement; In 94% of trials, it evoked a lateral movement (41%) or an arm curl (53%). On the other hand, Proximal and Cord stimulations did not induce a consistent response: Proximal stimulation induced lateral movement (25%), arm curl (29%), and in 46% of trial no movement was evoked; Cord stimulation induced lateral movement (31%), arm curl (31%), and in 38% of trial no movement was evoked.

**Figure 5.**
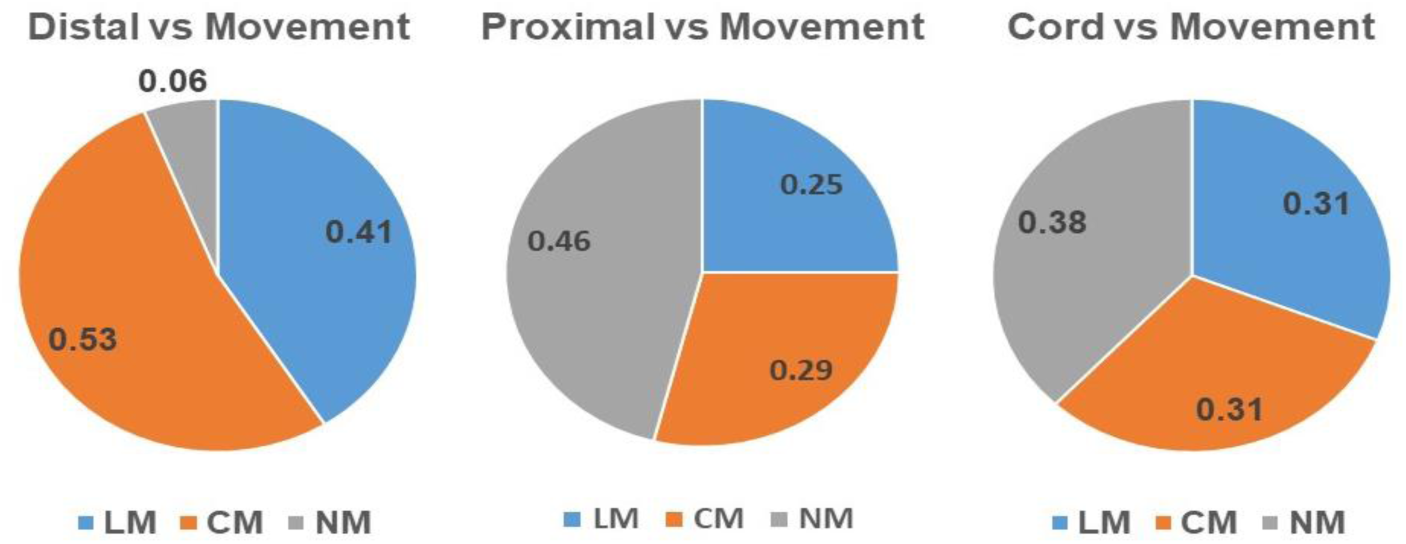
The stimulation type and movement response. (A) The probability of evoking each movement to occur after a proximal stimulation. (B) The probability of evoking each movement to occur after a Distal stimulation. (C) The probability of evoking each movement to occur after a Cord stimulation. This reveals that distal stimulation was very likely to induce a movement, whereas cord and proximal had a more even distribution of responses. (No movement (MN), movement with an arm curl (CM), lateral movement (LM).

The results showed that the number of units in the first 100ms post-stimulation can predict the movement response. The predictive probability of longer period of units to inform on movement response was examined. The total number of spikes in the first 500ms after stimulation was calculated for each of the 16 channels. The average number of units in the first 500ms for movement MN was 591±33.2, LM resulted in 691±40 units, and CM resulted in 706±54.7 spikes (average±SEM). An ANOVA analysis showed that there wasn’t a significant difference between the number of units to the movement response (F(2.00, 18.56)=0.39, p=0.68). In addition, there wasn’t a significant difference between the stimulation type and the number of units (F(2.00, 64.22)=0.45, p=0.64).

### Computational modelling of electrophysiology responses

#### Feature extraction

Temporal and stimulation features were extracted from the electrophysiology signals, using sample windows of 3s long, binned into 100ms, and summing the number of units in each window. This process resulted in a dataset with 30 bins that we treated as discrete features. To encode the stimulation information, categorical features were added using OHE technique to convert the categorical data into a numerical format. Two different datasets of 275 traces were created: a Multi-Class dataset where each sample was labelled 0 (no movement; 74 samples), 1 (movement; 96 samples), and 2 (movement with arm curl; 105 samples); and a Binary dataset where samples labelled 1 and 2 were combined into a single movement class (1, consisting of 201 samples), and the 0 class (74 samples). The distribution of the samples in the multi-class dataset was found to be balanced in the number of trials in each class. However, the distribution of the samples in the binary dataset showed a slight imbalance as it consisted of more samples in movement 1.

Next, we created a dataset by extracting temporal and stimulation features from the electrophysiology signals which contained 34 features: 33 features were predictors, and one feature was the categorical movement response (Target). Among 33 predictors, 30 predictors were discrete features, and the remaining three predictors were OHE features derived from the stimulation location: ANC stimulation, and stimulations located in the distal part and proximal part of the arm.

A Cramer’s V analysis was computed to understand the impact of binary features as shown in **Table 1**. The stimulation type feature was encoded using OHE technique. Then, Cramer’s V was computed between each binary feature and the target feature, reflecting the degree of association. Significantly, for both binary and multi-class targets, the Tactile, distal part binary feature is highly associated with the target (give me a number), indicating that the tactile, distal part binary feature is more useful for predicting the target features. The relatively low values for the ANC binary feature suggest it might be less useful for predicting the target features.

**Table 1.** Association strengths between binary features and target outcomes using Cramer’s V. The Cramer’s V analysis demonstrates the varying strengths of association, between the stimulation location features and the movement outcome. Results demonstrate that the tactile, distal location of stimulation feature, had higher association for both binary-class and multi-class outcomes, compared to ANC and tactile, proximal part features.

| Categorical Variables | Binary Target<br>(Cramér's V Value) | Multi-Class Target<br>(Cramér's V Value) |
| --- | --- | --- |
| Stimulation: ANC | 0.175 | 0.177 |
| Stimulation: Tactile, distal part | 0.435 | 0.500 |
| Stimulation: Tactile, proximal part | 0.287 | 0.373 |

Mutual information is a statistical method measuring the amount of dependence between two random variables ^39^, and **Figure 6** depicts the scores for both binary-class and multi-class.

**Figure 6:**
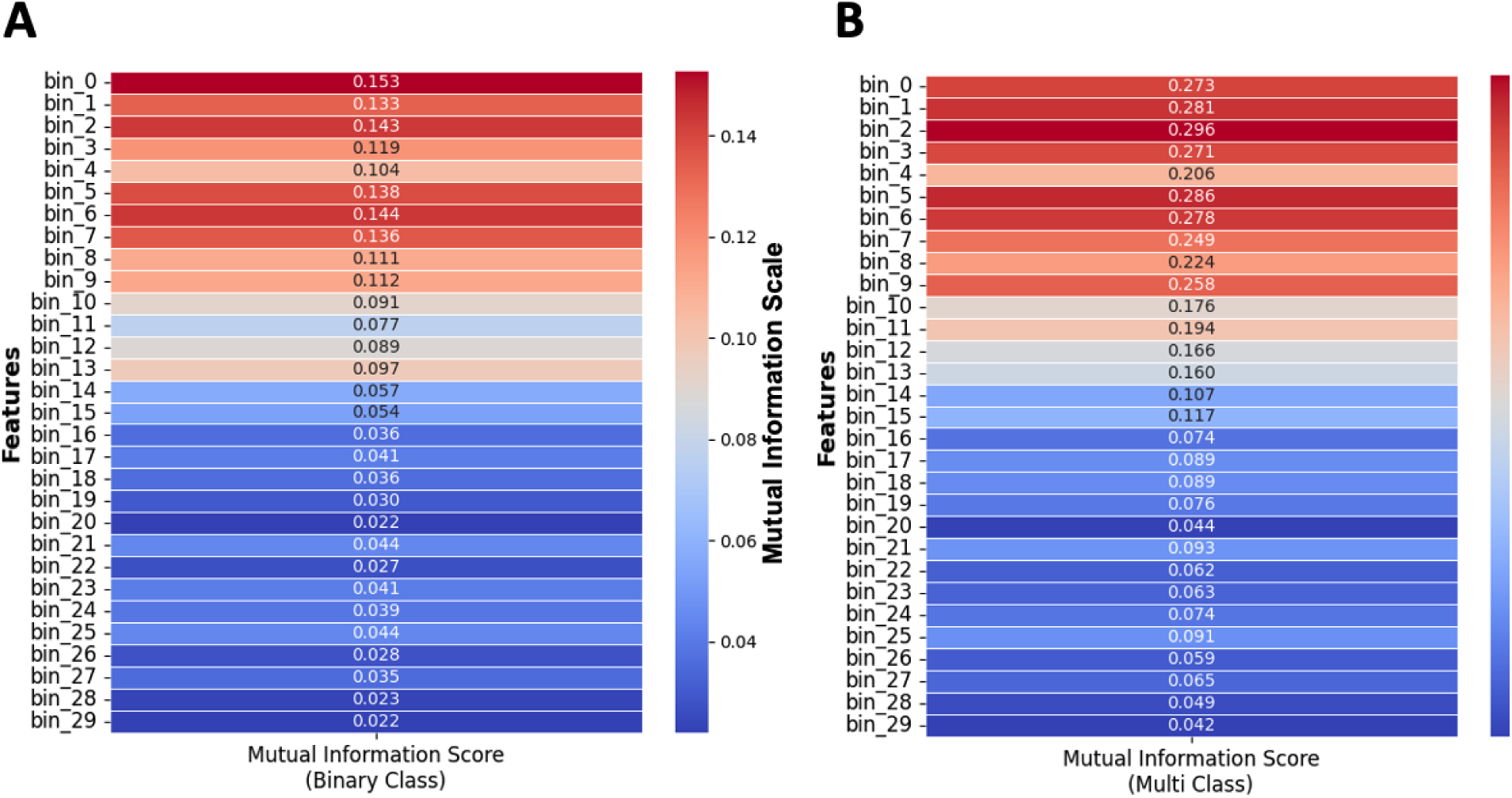
Mutual information analysis for understanding feature importance in binary and multi-class classification. (A) Mutual Information Scores between 30 binned discrete features, and Binary-Class scenario and (B) Mutual Information Scores between 30 binned discrete features, and Multi-Class scenario. Color bar indicates the mutual information score for a feature. This analysis demonstrates that the initial several hundred milliseconds after stimulation carry significant information about the target.

To identify any possible trends between the Mutual Information Scores of 30 binned, discrete input features and the output target, we performed a line fitting to these scores. This was done to further understand any existing linear trends. The R^2^ score for both Binary-Class and Multi-Class fitted lines were 0.85 and 0.87, respectively, and the slopes being -0.0047 and - 0.0095, respectively. These negative slopes that are also evident in **Figure 7** indicates that the significant dependence of the target decreases with time post-stimulation.

**Figure 7:**
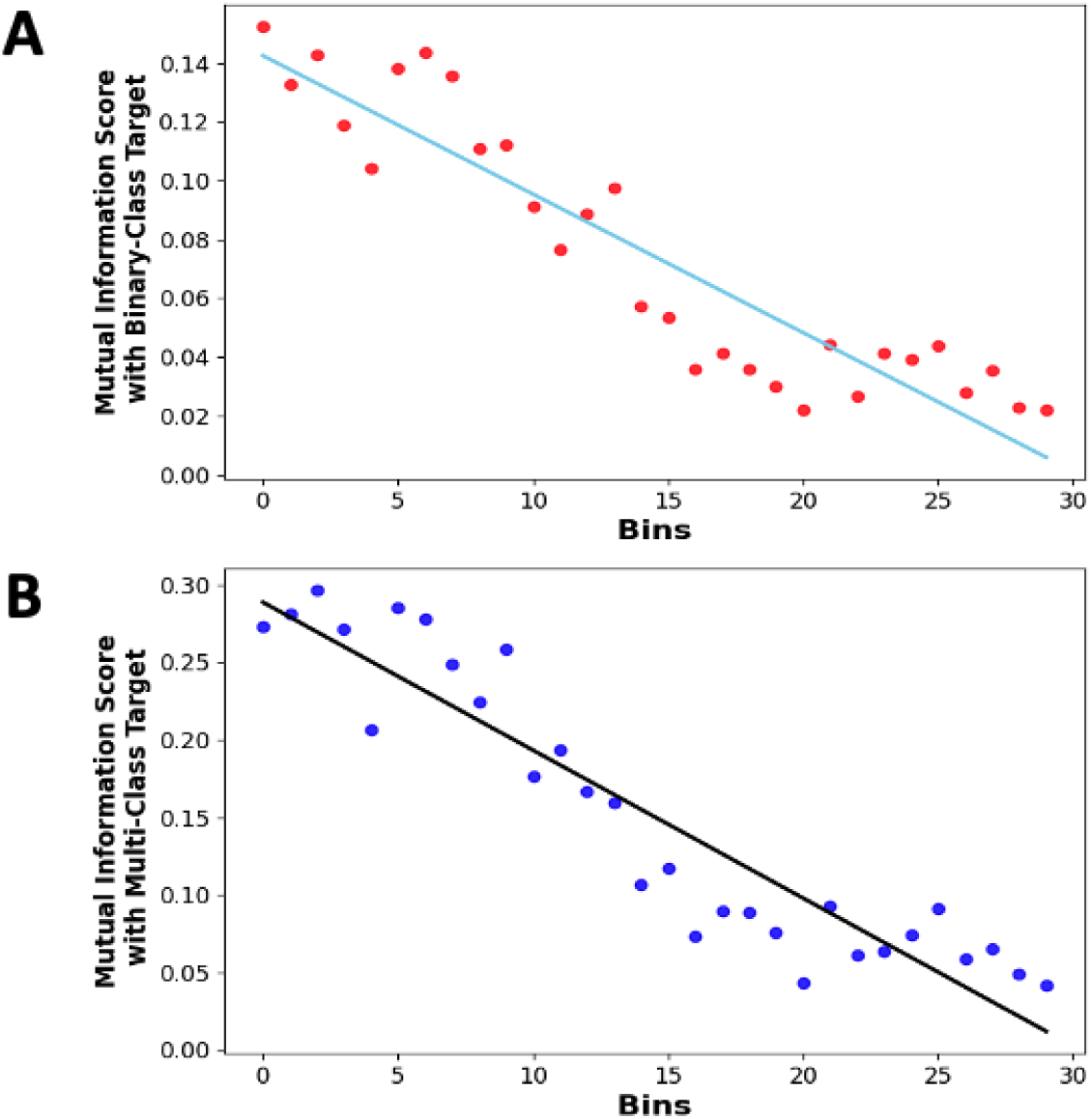
Mutual Information analysis for understanding feature significance in binary and multi-class classification. (A) Linear Trend analysis on Mutual Information Scores of 30 binned discrete features with the Binary-Class target, and (B) Linear Trend Analysis on Mutual Information Scores of 30 binned discrete features with the Multi-Class target. Each point indicates the mutual information score for a binned discrete input feature with target, and a fitted trend line showing the overall trend. The downward trends in both plots highlight that the significant dependence of the target on these features decreases over time, with the dependency diminishing as time approaches the end of the stimulation period (3000ms). This analysis aids in understanding which features are most influential in predicting the target in both binary and multi-class scenarios.

The feature importance analysis shown in **Figure 8** suggests that several input features are uniquely positioned to infer octopus arm movement, particularly within the first 100ms period of the electrophysiology response.

**Figure 8:**
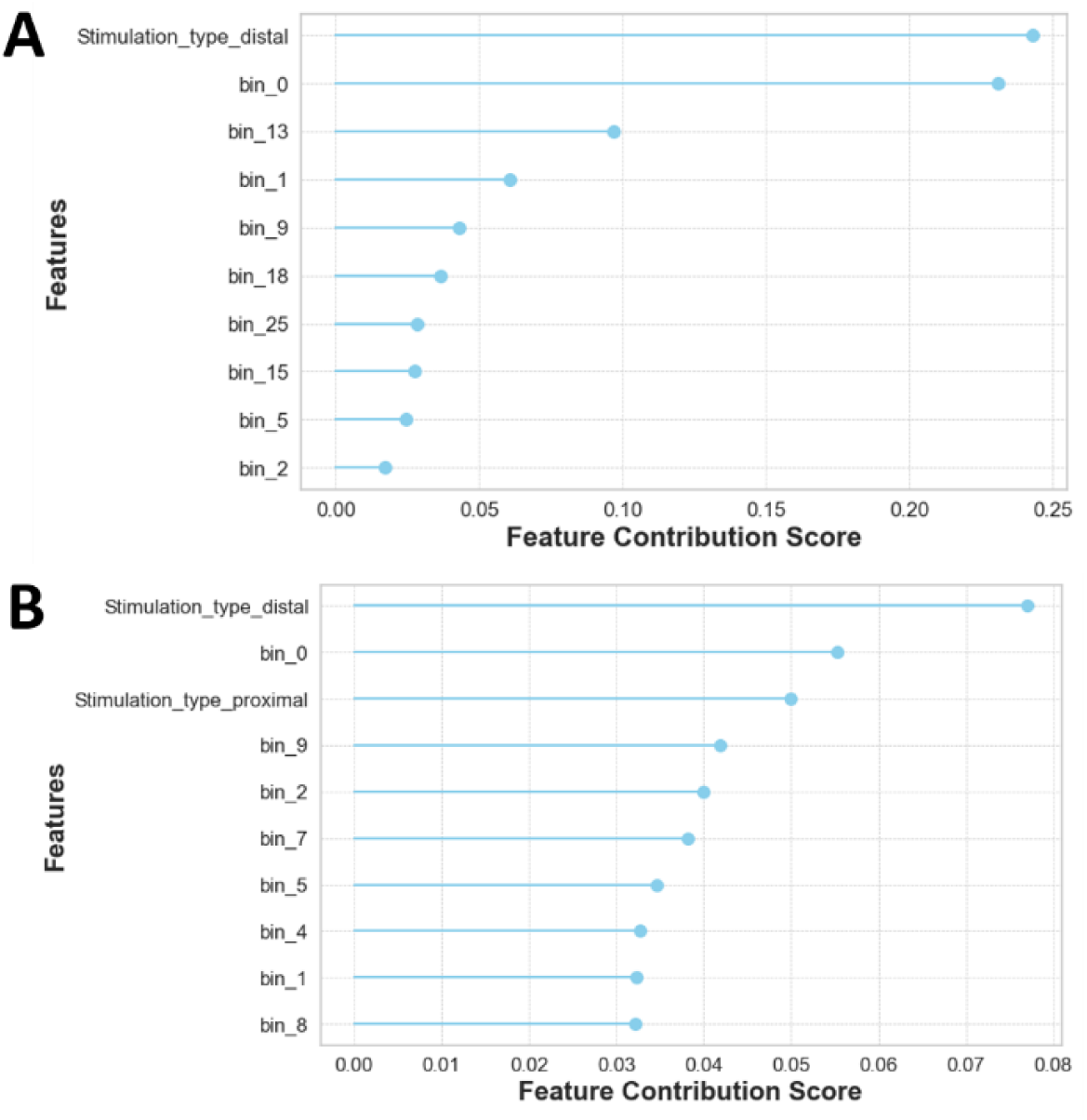
Decoding the top 10 Non-Linear dynamics via feature importance analysis. This analysis demonstrates the influence of different input features in model decision. It shows that for both (A) binary-class and (B) multi-class the type of stimulation and the timing of the feature is crucial.

#### Computational models that predict movement

The dataset was segregated by utilizing the 80/20 Split method, where 80% (220 samples) of the data was used for cross-validating of the model, and 20% (55 samples) of the data was used for testing the finalized model. The Stratified k-Fold Cross-validation technique was applied with K of 10 folds.

The accuracies of 16 different classifying techniques are reported in **Table 2**. These average performance scores returned from our cross-validation tests. Additionally, despite the class imbalance in the 100 ms binary dataset, with class 1 consisting of 201 samples and class 0 consisting of 74 samples, the model still appears to be performant and able to handle the imbalance effectively based on the evaluation of F1 score results from cross-validation (**Table 2**), and test dataset (**Table 3**). The best model binary-class dataset was the Gradient Boosting Classifier, which achieved 88.64% accuracy. The best model for the multi-class dataset was the Extra Trees Classifier, which achieved an accuracy of 75.45%. These models also showed the highest F1 scores, describing the harmonic mean of precision and recall, which reflects the model’s accuracy and consistency in identifying the correct category.

**Table 2:** Comparative analysis of Machine Learning classifier performance: Cross-validation metrics for binary-class and multi-class tasks. Eighty percent of the electrophysiology dataset was used for cross-validation. The most accurate models for prediction of movement for both multi-class and binary-class are represented with bold text.

| Model | Binary-Class |  |  |  | Multi-Class |  |  |  |
| --- | --- | --- | --- | --- | --- | --- | --- | --- |
|  | Accuracy | Recall | Precision | F1 | Accuracy | Recall | Precision | F1 |
| 1. Ada Boost Classifier | 81.36 | 88.12 | 87.36 | 87.24 | 60.45 | 60.18 | 64.6 | 60.32 |
| 2. CatBoost Classifier | 85.91 | 94.38 | 87.81 | 90.73 | 75 | **75.7** | **77.49** | **74.53** |
| 3. Decision Tree Classifier | 78.64 | 82.65 | 88.1 | 84.86 | 58.64 | 59.1 | 61.34 | 58.6 |
| 4. Dummy Classifier | 73.18 | 100 | 73.18 | 84.51 | 38.18 | 33.33 | 12.73 | 18.41 |
| **5. Extra Trees Classifier** | 83.18 | 92.54 | 86.12 | 88.98 | **75.45** | **75.58** | **77.32** | **74.47** |
| 6. Extreme Gradient Boosting | 82.27 | 89.38 | 87.74 | 88.08 | 69.09 | 69.58 | 70.97 | 67.83 |
| **7. Gradient Boosting Classifier** | **88.64** | **93.12** | **92.12** | **92.34** | 72.73 | 72.87 | 74.45 | 72.23 |
| 8. K Neighbors Classifier | 78.18 | 88.86 | 83 | 85.62 | 56.82 | 57.8 | 60.22 | 56.12 |
| 9. Light Gradient Boosting Machine | 86.36 | 91.91 | 90.33 | 90.78 | 70 | 70.05 | 71.01 | 69.11 |
| 10. Linear Discriminant Analysis | 80.45 | 86.36 | 87.32 | 86.53 | 66.82 | 67.83 | 69.17 | 66.1 |
| 11. Logistic Regression | 80.91 | 87.65 | 86.58 | 86.91 | 61.36 | 61.87 | 62.74 | 60.76 |
| 12. Naive Bayes | 78.18 | 78.2 | 91.25 | 83.69 | 43.64 | 42.51 | 47.83 | 40.94 |
| 13. Quadratic Discriminant Analysis | 80.45 | 97.54 | 80.23 | 87.98 | 67.73 | 65.53 | 74.27 | 65.51 |
| 14. Random Forest Classifier | 83.64 | 93.75 | 85.82 | 89.29 | 72.73 | 72.38 | 74.3 | 71.72 |
| 15. Ridge Classifier | 80.45 | 86.99 | 86.74 | 86.59 | 63.64 | 64.15 | 66.32 | 62.58 |
| 16. SVM - Linear Kernel | 71.36 | 79.78 | 84.03 | 78.74 | 50.45 | 51.16 | 51.03 | 47.34 |

**Table 3.** Evaluation of Machine Learning Classifiers on Test Data: Performance metrics across Binary and Multiclass classification. The two models outperformed during the cross-validation and were tested on 20% of the test data. These results demonstrate that the models can predict the movement with high accuracy even when it was tested on new electrophysiology data.

| Model | Accuracy (Binary-Class) | Recall (Binary-Class) | Precision (Binary-Class) | F1 (Binary-Class) | Accuracy (Multi-Class) | Recall (Multi-Class) | Precision (Multi-Class) | F1 (Multi-Class) |
| --- | --- | --- | --- | --- | --- | --- | --- | --- |
| 1. Extra Trees Classifier | -- | -- | -- | -- | 72.73 | 72.65 | 74.11 | 73.22 |
| 2. Gradient Boosting Classifier | 83.64 | 97.50 | 82.98 | 89.66 | -- | -- | -- | -- |

The performance metrics for the test data are presented in **Table 3**. The Gradient Boosting Classifier, identified as the best model through cross-validation on the Binary-Class dataset, was tested on 20% of previously unseen data and achieved an accuracy of 83.64%. This compares to the 88.64% accuracy obtained with the stratified k-Fold evaluation on the remaining 80% of the data. Similarly, the Extra Trees Classifier, which was the best model from cross-validation on a Multi-class dataset, was tested on 20% of unseen data and reached an accuracy of 72.73%, versus the 75.45% accuracy achieved with the stratified k-Fold evaluation on the remaining 80% of the data

This analysis indicated that the Gradient Boosting Classifier could predict with 88.64% accuracy the movement in datasets composed of 30 bins of 100ms each. The accuracy of movement prediction using 50ms bins was also tested and compared to the 100ms dataset. This dataset consisted of 60 continuous binned features, the categorical features, and the Target features. The results demonstrated that higher accuracy in movement prediction could be achieved with 100ms compared to the 50ms dataset, as shown in **Table 4**.

**Table 4.**
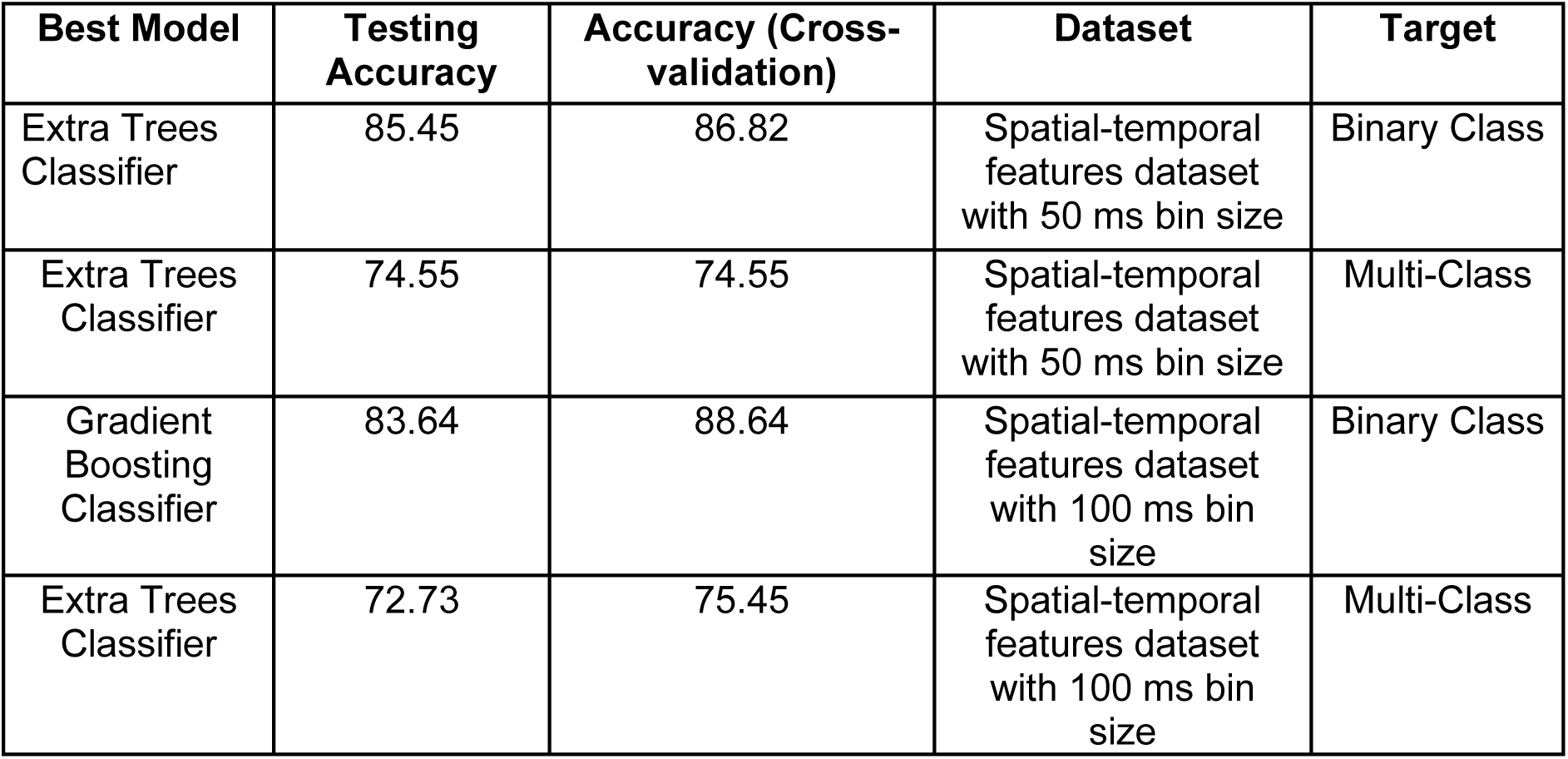
Comparison of models across time frames. The outperforming models were tested on electrophysiology signals binned into 100ms and 50ms features. Comparing the accuracy of the results from the cross-validation shows that 100ms features have led to 1.82% higher accuracy in Binary-class and 0.9% higher accuracy in the Multi-class compared to the 50ms features.

The confusion matrix (**Figure 9**) is a method that allows computing a machine learning model accuracy, precision, recall, and overall ability to correctly classify instances, providing a detailed view of the types and frequencies of classification errors. These results suggest that the Gradient Boosting Classifier and the Extra Trees Classifier could predict type of movement with high accuracy.

**Figure 9:**
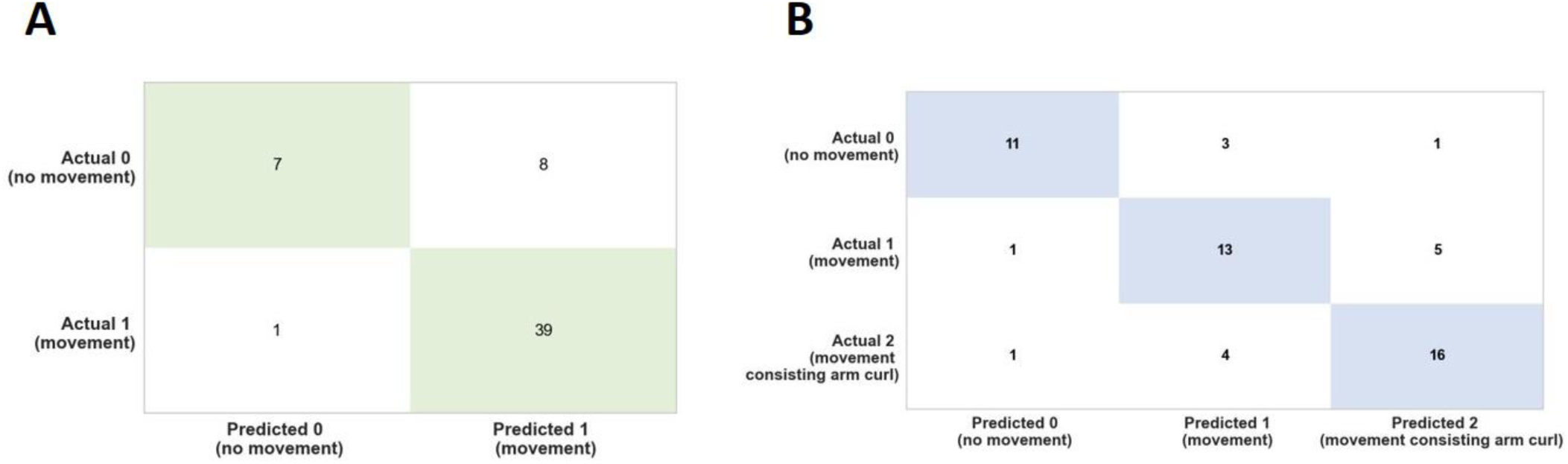
Confusion Matrix analysis for binary and multi-class movement predictions. (A) Binary-class and (B) Multi-class confusion matrix analysis showing the correct and incorrect predictions of the type of movement based on the test dataset which is 20% of the entire data. The green and blue indicate the correct predictions. This analysis suggests the high accuracy of the models in predicting the type of movement.

### Computational models of movement to decode stimulation

We next considered a finer-grained analysis of the movements evoked from stimulation of the arm to determine relevant kinematic features beyond the 0, 1, and 2 movement class labels manually applied. Different stimulation locations elicited distinct behavioral responses in octopus arm movement. We examined the distribution of each kinematic metric previously defined, using both keypoint-derived features and proSVD identified bases. **Figure 10** shows one example of how movement evoked from stimulation at the cord, versus at the distal or proximal regions (PD), has a significantly different distribution of the maximum angular velocity across each time period post-stimulation (see **Appendix** for complete table). Initial analyses considered the distal and proximal regions separately, but found no significant differences, and so all future analyses considered them as a single combined group.

**Figure 10:**
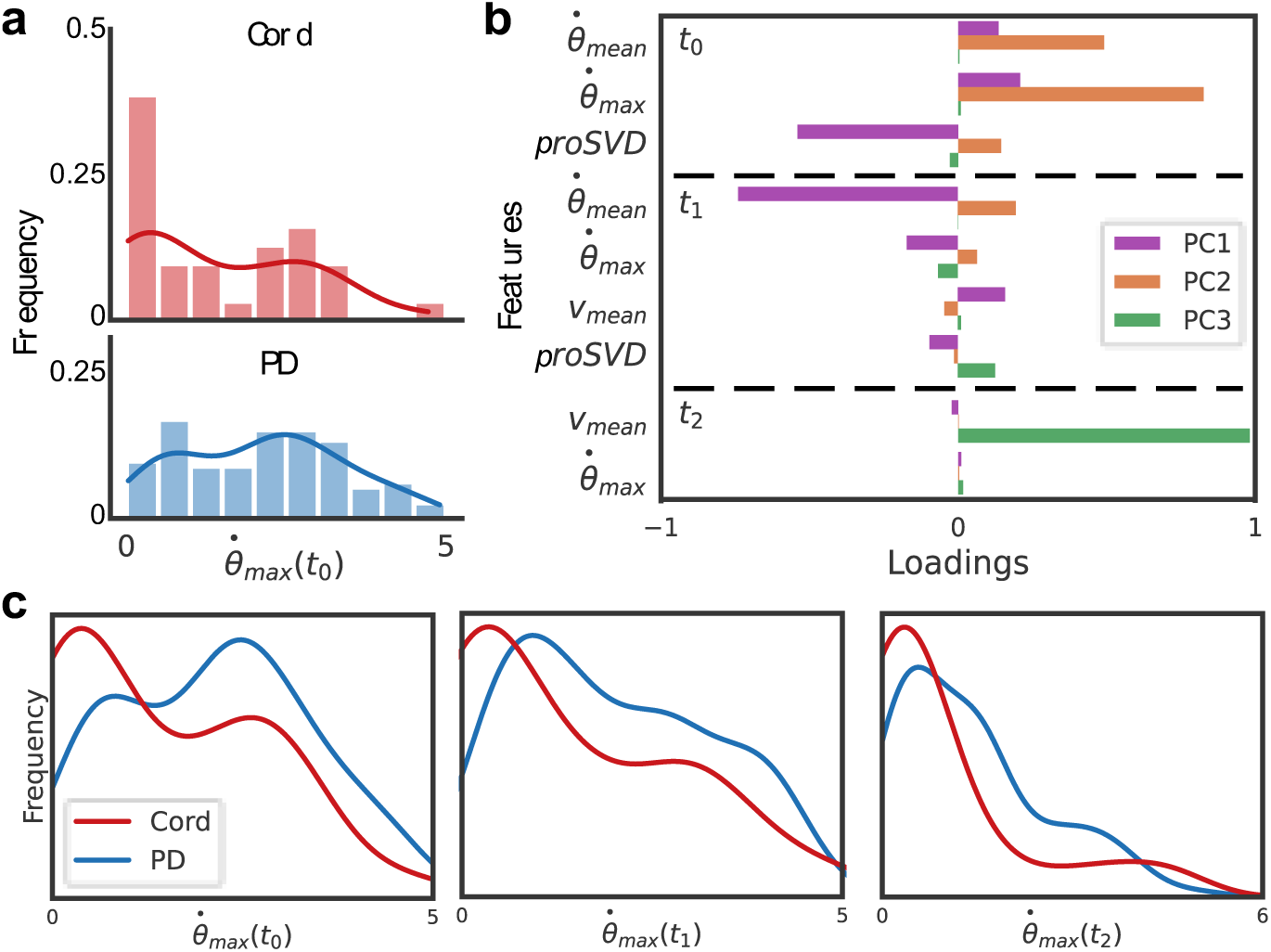
Different motion features evoked by stimulus location. (A) Histograms and overlaid kernel density estimation plots for the extracted maximum angular velocity in t0 for each video during (top) cord or (bottom) proximal or distal (PD) stimulation. (B) A principal component analysis was performed for all extracted behavior features. Selected features are shown here with higher loadings across the first three principal components (PC). The mean and maximum angular velocities and proSVD features across the first two time periods comprise much of the first two PCs, with the distal velocity in the last time period having the highest loading onto the third PC. (C) The distributions of this metric, maximum angular velocity, are significantly different across all post-stimulation time windows (for t0, t1, t2; p=0.014, p=0.001, p=0.004, respectively, determined by a two-sample KS test).

Principal component analysis was conducted on all features across all time periods to identify which features might be playing the largest role (selected features shown in **Figure 10b**). Immediately following stimulation, the angular speed of the distal part and the proSVD bases were the most significant kinematic features, with translation speed contributing more significantly in the later time periods. We found that the features with the highest loadings in PCA space also tended to produce significantly different feature distributions between stimulation types.

The above analyses all collapsed movement into three time periods, potentially missing relevant signals at finer time resolution. To characterize more complex behavior, and to establish real-time methods for future closed-loop work, we looked at what metrics and analyses we could do in the streaming setting, as fast as data could be collected. DLC live was able to generate keypoint inferences at rates of ∼10-20 ms per image, or around 100 frames per second on average (**Figure 11a**). proSVD could be run at extremely high frame rates, and may capture finer motion features than those that could be identified using keypoint tracking. We found that proSVD features showed some differences between mechanical and electrical stimulation types as a function of time post-stimulation (**Figure 11b**), unlike earlier keypoint metrics. While less interpretable, we anticipate that including these types of unsupervised features alongside user-defined keypoints will produce the quantification needed to fully characterize the rich and complex behavior in the octopus repertoire.

**Figure 11:**
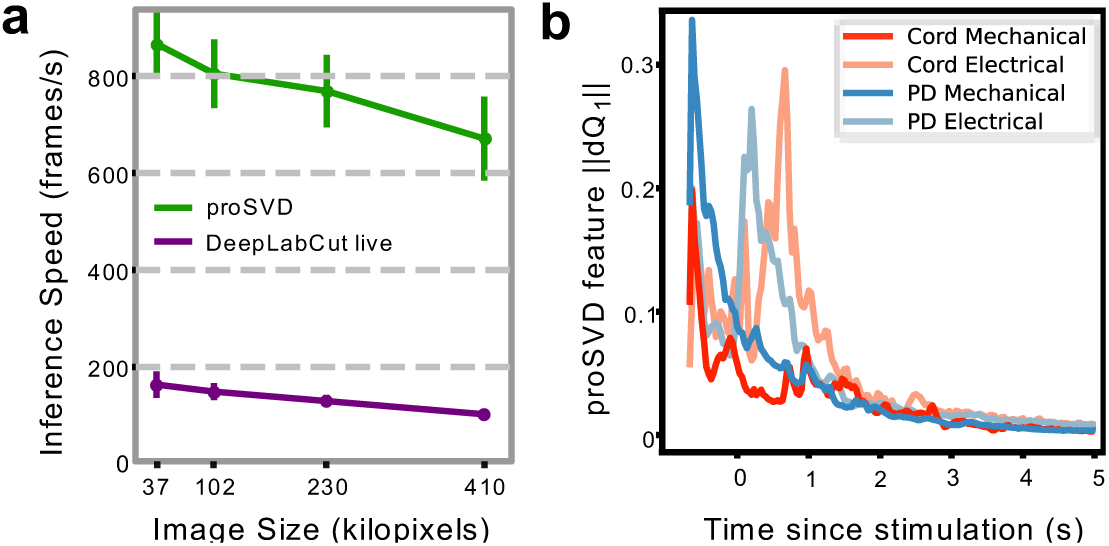
Real-time inference and time-varying features. a, Inference speeds as a function of video resolution for both DeepLabCut live and proSVD during streaming analyses for different downsampled video resolutions. Error bars denote standard deviation (N=10). b, The L2-norm of the derivative of the first feature identified by proSVD plotted as a function of time post-stimulation, averaged across all trials. The low-dimensional representation of the evoked motion shows some different temporal features for each stimulus conditions.

## Discussion

The octopus’s extraordinary anatomy and physiology makes it especially attractive to uncover sensorimotor circuits that orchestrate behavior. The octopus’s nervous system is highly distributed, and much of the neural circuitry coordinating these behaviors is organized within the arms where it can monitor immediate, complex environmental feedback and adapt the arms’ movement accordingly ^7,8,40,41^. Identifying and understanding the neural signals that drive complicated motor output, such as those found in the octopus, are essential for the future development of rapidly-adapting and ultimately more human-like prosthetic arms ^42,43^.

Here we describe high-temporal and spatial resolution results of single and multi-unit data, obtained from a detached, behaving octopus’s arm. The results show that the number of spikes occurring within the first 100ms after stimulation can predict the movement response, whereas the greatest number of spikes were associated with lateral movement. Stimulation location was also a significant variable: The greatest number of spikes in the first 100ms occurred in response to proximal stimulation, but distal stimulation evoked the greatest change in spike response over time. These results indicate that spike analysis can reveal fundamental principles of motor behavior.

While the single and multiunit analysis by itself consisted of important information of behavior, we tested if using ML could further identify important features of octopus motor behavior. The models that were built on the ML results indicate that it was possible to predict whether an arm movement occurred with 88.64% confidence, and it was possible to predict with 75.45% confidence if this was a lateral arm movement or an arm movement with a curl. These levels of prediction values are in agreement with the range of reported accuracies in arm reaching predictions based on large-scale neural activity in monkey’s motor cortex ^44,45^ and in humans ^46^. The accuracy of machine learning algorithms in predicting arm movements can vary depending on several factors, including the specific algorithm used, the quality and quantity of input data, the level of noise, and the complexity of the movement being predicted.

Consistent with the unit analysis results, the feature importance analysis showed that the first 100 ms period of the electrophysiology response is the strongest feature that predicts the type of movement. On top of the electrophysiology signals, it was important to include the stimulation location in the model, specifically the peripheral stimulation to the distal part of the arm. The importance of the distal location is consistent with other evidence showing that that distal stimulus evokes a response with the shortest latency and the highest amplitude compared to proximal stimuli or stimulus of the ANC ^13^. The motion analysis results showed that the same stimulation in the octopus does not always produce the same behavior, supporting that the movement of the detached arm is a complex movement triggered by central and peripheral neural circuits ^10,11,47^.

We have recently used a set of five reflective markers that were adhered to the octopus skin, in order to describe and quantify the overall posture of an awake, swimming octopus ^48^. Three postures that were defined as straight, simple bending, and complex bending, and were analyzed in 3D using curvature and plane orientation methods. The results showed that this novel kinematics approach was successful in understanding octopus posture. However, this approach is limited by the number of markers that could be attached to the octopus’s arm and the physical constrains they might induce.

Nevertheless, state-of-the-art computer vision and machine learning tools could provide quantification of kinematic features based on video recordings alone ^49–51^. Deep learning and other machine learning methods could also learn features that human eyes do not see, but may be significantly correlated with neural firing or stimulation patterns ^52–54^. Employing transfer learning, deep neural networks, and dimension reduction as described here, we aimed to gain real-time insights into octopus arm movements and how their motor circuitry produces rich movement types.

There are limitations in current methods for accurately locating points of interest. In the octopus arm videos considered here, several display minimal to no movement except for mechanical adjustments made by the experimenter. The predominance of single-instance movement in most videos, with shorter clips, effectively limits our dataset for training the DLC model on complicated movements. Additionally, the non-planar motion of the arm at times poses a challenge for accurate tracking, requiring new tracking strategies to capture the full complexity of octopus arm movements.

Our selection of kinematic parameters was inspired by a study on locomotion using zebrafish larvae ^26^, but their effectiveness for the octopus arm is not straightforward due to distinct ethology and experimental conditions. Unlike zebrafish, the octopus arm lacks a zero-angle “tail” at rest. Zebrafish data were made positionally consistent through affine transformations and background removal, a step not directly applicable to octopus arm data due to sample size limitations, hindering common clustering techniques. Assessing the effectiveness of unsupervised clustering to identify key features also proves challenging in the absence of ground truth labels to gauge cluster accuracy and precision. Overall, improvements could be achieved through higher resolution videos, camera stability without flickering, incorporation of multiple stable camera angles, precise manual annotation, and a larger volume of data.

The experiment aimed to uncover insights into octopus arm movements through meticulous video analysis. The electrical and mechanical stimuli induced diverse responses in the octopus arms, ranging from no movement to complex arm curls. The kinematic analysis and feature extraction provided valuable quantitative data, shedding light on key aspects of the arm’s motion, such as angles, angular speed, and absolute speed. These outcomes can collectively contribute to a deeper understanding of octopus arm behavior and provide a foundation for further investigations into motor control and neural circuitry. Beyond enhancing our understanding of neural circuits, this work has potential implications in brain-machine interfaces and prosthetics, enabling the development of sophisticated systems that replicate natural movement with precision and fine temporal resolution.

## Acknowledgments

The authors would like to acknowledge financial support from the NIH: UF1NS115817; R01NS098231.

## Author contributions

Conceptualization: G.P.; Methodology: N.G., S.S, J.R., C.C, A.D., G.P.; Analysis; N.G., G.P. S.S, R.R., A.D.; Writing-Original Draft: N.G., S.S., R.R., A.D., G.P.; Writing-Review and Editing: N.G., S.S., J.R., R.R., C.C., A.D., G.P.

## Data availability

The data that support the findings of this study are available from the corresponding author upon reasonable request. The codes will be shared through Github.

## Appendix

| K-S Significance Test on PD (Proximal-Distal v/s Cord) |  |  |  |  |
| --- | --- | --- | --- | --- |
| Time period | Statistic | Feature | p-value | distribution |
| t_0 | Mean | Angular Speed | 0.010 | different |
|  | Max |  | 0.014 | different |
|  | Mean | Distal Speed | 0.257 | same |
|  | Max |  | 0.016 | different |
|  | Max | ProSVD feature | 0.753 | same |
| t_1 | Mean | Angular Speed | 0.001 | different |
|  | Max |  | 0.001 | different |
|  | Mean | Distal Speed | 0.029 | different |
|  | Max |  | 0.012 | different |
|  | Max | ProSVD feature | 0.107 | same |
| t_2 | Mean | Angular Speed | 0.003 | different |
|  | Max |  | 0.004 | different |
|  | Mean | Distal Speed | 0.085 | same |
|  | Max |  | 0.051 | same |
|  | Max | ProSVD feature | 0.012 | different |

